# Skyhawk: An Artificial Neural Network-based discriminator for reviewing clinically significant genomic variants

**DOI:** 10.1101/311985

**Authors:** Ruibang Luo, Tak-Wah Lam, Michael C. Schatz

## Abstract

**Motivation:** Many rare diseases and cancers are fundamentally diseases of the genome. In the past several years, genome sequencing has become one of the most important tools in clinical practice for rare disease diagnosis and targeted cancer therapy. However, variant interpretation remains the bottleneck as is not yet automated and may take a specialist several hours of work per patient. On average, one-fifth of this time is spent on visually confirming the authenticity of the candidate variants.

**Results:** We developed Skyhawk, an artificial neural network-based discriminator that mimics the process of expert review on clinically significant genomics variants. Skyhawk runs in less than one minute to review ten thousand variants, and about 30 minutes to review all variants in a typical whole-genome sequencing sample. Among the false positive singletons identified by GATK HaplotypeCaller, UnifiedGenotyper and 16GT in the HG005 GIAB sample, 79.7% were rejected by Skyhawk. Worked on the Variants with Unknown Significance (VUS), Skyhawk marked most of the false positive variants for manual review and most of the true positive variants no need for review.

**Availability:** Skyhawk is easy to use and freely available at https://github.com/aquaskyline/Skyhawk

## Introduction

The dramatic reduction in the cost of whole genome, exome and amplicon sequencing has allowed these technologies to be increasingly accessible for genetic testing, opening the door to broad applications in Mendelian disorders, cancer diagnosis and personalized medicine [1]. However, sequencing data include both systematic and random errors that hinder any of the current variant identification algorithms from working perfectly. Even using state-of-the-art approaches, typically 1-3% of the candidate variants are false positives with Illumina sequencing [2]. With the help of a genome browser such as IGV [3], or web applications such as VIPER [4], a specialist can visually inspect a graphical layout of the read alignments to assess supporting and contradicting evidence to make an arbitration. Though necessary, this is a tedious and fallible procedure because of three major drawbacks. 1) It is time-consuming and empirical studies report it requires about one minute per variant, sometimes summing up to a few hours per patient [5]. 2) It is tedious, not infallible, and even experienced genetic-specialists might draw different conclusions for a candidate variant with limited or contradicting evidence. 3) There is no agreed standard between genetic-specialists to judge various types of variants, including SNPs (Single Nucleotide Polymorphisms) and Indels. A specialist might be more stringent on SNPs because there are more clinical assertions and fewer candidate SNPs will be less likely to get contradicting medical conclusions, whereas another specialist might be more demanding on indels because they are rarer and harder to be identified.

An efficient, accurate and consistent computational method is strongly needed that automates assessing the candidate variants as they would be visually validated. Importantly, the new validation method needs to be orthogonal, i.e., independent of the algorithms used to identify the candidate variants. The new validation method also needs to capture the complex non-linear relationship between read alignments and the authenticity of a variant from a limited amount of labeled training data. Variant validation is a task with a different nature from variant filtration. Our target is to indicate the need of a variant being manually reviewed, as opposed to a hard filter that removes a variant from consideration. To achieve our target, failing to flag a false positive variant for review is less favorable than flagging a true variant for manual review, i.e., as a validation method, the precision must be maximized, and false positives must be minimized. Consequently, instead of using hand-coded models or rule-based learning, a more powerful and agnostic machine learning approach such as an Artificial Neural Network (ANN) is needed.

## Implementation

We implemented Skyhawk, a computational discriminator that is fast and accurate for validating candidate variants in clinical practice. Skyhawk mimics how a human visually identifies genomic features comprising a variant and decides whether the evidence supports or contradicts the sequencing read alignments. To reach this goal, we repurposed the network architecture we developed in a previous study named Clairvoyante [6]. The multi-task ANN was designed for variant calling in Single Molecule Sequencing, and the method is orthogonal to traditional variant callers using algorithms such as Bayesian or local-assembly. In Skyhawk, we used a repurposed network to generate a probability of each possible option for multiple categories including 1) variant type, 2) alternative allele, 3) zygosity, and 4) indel-length. We then compare a candidate variant to Skyhawk’s prediction on each category. Skyhawk will agree with a variant if all categories are matched but will reject and provide possible corrections if any category is unmatched. We have provided pre-trained models for Skyhawk on GitHub trained using the known variants and Illumina data of multiple human genomes, including sequencing libraries prepared by either the PCR or the PCR-free protocol. With a trained model, Skyhawk accepts a VCF input with candidate SNPs and Indels, and a BAM input with read alignments. Skyhawk outputs a judgment and a quality score on how confident the judgment was made for each candidate variant. Skyhawk was implemented in Python and Tensorflow and has been carefully tuned to maximize its speed.

## Results

Using four deeply Illumina sequenced genomes (HG001, HG002, HG003, and HG004) with 13.5M known truth variants from the Genome In A Bottle (GIAB) project [2], we trained Skyhawk to recognize how the truth variants are different from another 20M non-variants we randomly sampled from the four genomes. The sample details and the commands used are in the **Supplementary Note**. For benchmarking and identifying the false positive variant calls, we used the known truth variants in HG005, which was not included in the model training. A false positive variant is defined as a variant called by a variant caller but cannot be found in the HG005 GIAB truth dataset and will be used for the subsequent analysis. We expect the false positive variants that are supported by only one variant caller, but not the other variant callers are very likely to be erroneous and should be marked for manual review (i.e., rejected by Skyhawk) [7]. Thus, we called variants using three different variant callers with different calculation models, including GATK HaplotypeCaller (HC) [8], GATK UnifiedGenotyper (UG) [8] and 16GT [9]. A Venn diagram of the variant set called by the three callers comprise seven different types of variant: 1) three types of singleton variant that have support from only one caller, 2) three types of doubleton variant that have support from two of the three callers, and 3) one type of tripleton variant that is supported by all three callers. Empirically, doubleton and especially tripleton variants are relatively less likely to be real false positives and should be less likely to be rejected by Skyhawk. Conversely, singletons called by only one caller are more likely to be genuine false positive and should be more likely to be rejected by Skyhawk. The results are shown in **Figure 1**. Only 18.64% of the tripleton variants were rejected while 79.70% of the singletons were rejected by Skyhawk. Those doubletons have an intermediate 45.11% rejected by Skyhawk. In the true positive variants, only 1,879/3,232,539 (0.058%) in HC, 43/2,902,052 (0.0014%) in UG, and 124/3,228,537 (0.0038%) in 16GT were rejected. By deducting the rejected variants from both the number of true positives and true negatives, the precision increased from 99.77% to 99.92% for HC, 99.50% to 99.58% for UG and 99.51% to 99.84% for 16GT.

**Figure 1.**
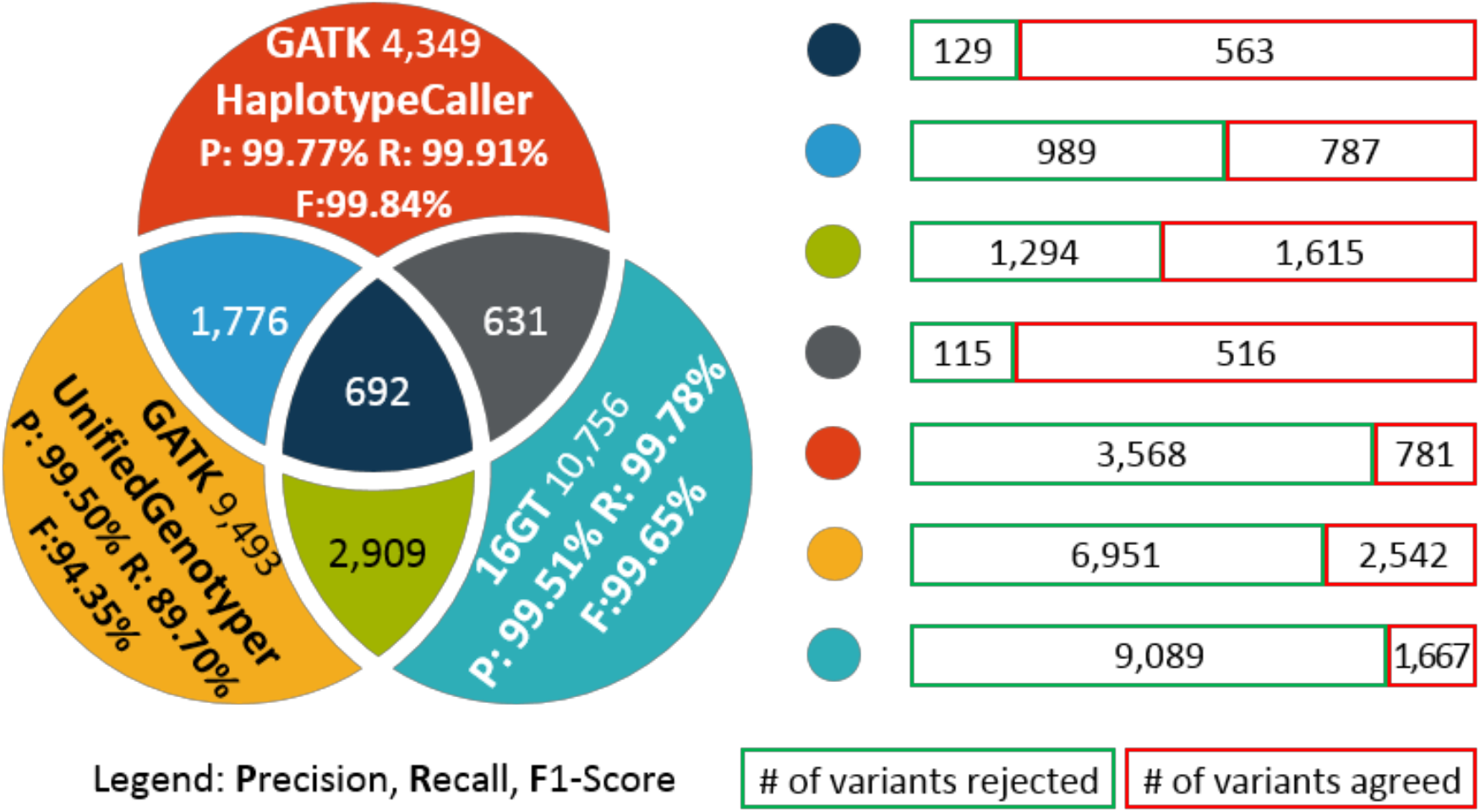
The variant calling results of GATK HaplotypeCaller, GATK UnifiedGenotyper, and 16GT. The Venn diagram on the left shows 1) the precision rate (P), recall rate (R) and f1-score (F) of each variant caller on all variants of the entire HG005 genome, and 2) the number of false positive variants produced by each variant caller. The bars on the right shows the number of false positive variants rejected or agreed by Skyhawk. The bar length is proportionate to the total number of false positive variants in that type.

Another experiment better mimics how medical doctors would use Skyhawk in clinical diagnosis. Instead of fully removing manual review, which is impossible in a stringent clinical context the emphasizes accountability, Skyhawk’s target is to help doctors to prioritize which variants should they invest efforts in further investigation and lab validation. In practice, those variants categorized as “Pathogenic” or “Likely Pathogenic” are rare and should be given priority [10], thus all these variants are preferred to be manually reviewed. “Benign”, and most of the time together with the “Likely Benign” category, suggest variants without much value in clinical diagnosis and therapy, thus not requiring manual review. The one category left, named Variant of Unknown Significance, or VUS, contains variants that are potentially impactful, and requires doctors to sort through them. The number of VUS is usually tens to even hundreds of time larger than the sum of other categories [11]. Thus, Skyhawk will benefit the clinical doctors if it can significantly decrease the number VUS to be manually reviewed. To assess the intended function, we firstly ran GATK HaplotypeCaller on the HG002 sample. In total about 5M variants were called. Then we annotated all variants using SeattleSeq version 151 (with dbSNP v151) [12]. We extracted those variants that are 1) not in dbSNP (RSID tag equals to 0) and, 2) are in a human gene (GL tag not empty). Finally, we ran Skyhawk on the extracted variants with a model trained on four samples including HG001, HG003, HG004, and HG005, and annotated the variants as either true positive (TP) or false positive (FP) against the HG002 GIAB truth dataset. Skyhawk performed as expected, and the results are shown in **Table 1**. For SNPs, 53.4% of the FPs are flagged for manual review, while only 0.3% of the TPs are flagged. For Indels, 78.3% of the FPs are flagged for manual review, while only 25.5% of the TPs are flagged. A higher rate of TP Indels is flagged for manual review because longer Indels are usually more error-prone and can lead to more several clinical consequences than SNPs, thus we required all Indels ≥4bp to be manually reviewed. Noteworthy, although an ideal percentage of FP being marked for manual review is 100%, it is not yet achievable because as mentioned in the previous paragraph, FP still have a chance to be an authentic variant especially when it is supported by multiple variant callers. Nevertheless, the trend of having significantly more FP variants marked for manual review than TP variants verified Skyhawk’s effectiveness.

**Table 1.** Skyhawk’s performance on Variants of Unknown Significance (VUS)

|  |  | PASS |  | CHECK |  |
| --- | --- | --- | --- | --- | --- |
|  |  | # | % | # | % |
| TP | SNP | 4,837 | 99.7% | 14 | 0.3% |
|  | Indel | 7,126 | 74.5% | 2,434 | 25.5% |
| FP | SNP | 117 | 46.6% | 134 | 53.4% |
|  | Indel | 41 | 21.7% | 148 | 78.3% |

## Discussion and Conclusions

Skyhawk aims to relieve users from a heavy manual review workload without compromising the accuracy. Instead of taking over the review of all variants, Skyhawk was configured to review only 1) SNPs with a single alternative allele, and 2) Indels ≤4bp. Skyhawk also outputs a quality score ranging from 0 to 999 to indicate how confident a judgment is. Among the false positive singletons, 27.46% of the judgments were with a quality score lower than 150. Reviewing these variants manually shows that these variants were often located in genome regions with homopolymer runs or very low depth. We suggest users to rely on Skyhawk only when the quality score of judgment is high and to manually review when the quality score falls below 150, or higher if the workload allows. Skyhawk requires less than a gigabyte of memory and less than a minute on one CPU core to review ten thousand variants, thus can be easily integrated into existing manual review workflows, such as VIPER [4] with minimal computational burden. Using 24 CPU cores, Skyhawk was able to review all five million whole genome sequencing variants of the HG002 sample in 30 minutes. Overall, Skyhawk greatly reduces the workload on reviewing variants, and we believe Skyhawk will immediately increase the productivity of genetic-specialists in clinical practice.

## Acknowledgment

R.L. was supported by the General Research Fund No. 27204518. T. L. was partially supported by Innovative and Technology Fund ITS/331/17FP from the Innovation and Technology Commission, HKSAR. This work was also supported, in part, by awards from the National Science Foundation (DBI-1350041) and the National Institutes of Health (R01-HG006677).

