## Supplementary Materials for "Skyhawk: An Artificial Neural Network-based discriminator for reviewing clinically significant genomic variants"

### Supplementary Note

#### Data Source

##### Truth Variants (Genome in a Bottle dataset version 3.3.2)

###### *HG001 (NA12878), GRCh38*

ftp://ftp-trace.ncbi.nlm.nih.gov/giab/ftp/release/NA12878_HG001/NISTv3.3.2/GRCh38

###### *HG001 (NA12878), GRCh37*

ftp://ftp-trace.ncbi.nlm.nih.gov/giab/ftp/release/NA12878_HG001/NISTv3.3.2/GRCh37

###### *HG002 (NA24385), GRCh38*

ftp://ftp-trace.ncbi.nlm.nih.gov/giab/ftp/release/AshkenazimTrio/HG002_NA24385_son/NISTv3.3.2/GRCh38

###### *HG002 (NA24385), GRCh37*

ftp://ftp-trace.ncbi.nlm.nih.gov/giab/ftp/release/AshkenazimTrio/HG002_NA24385_son/NISTv3.3.2/GRCh37

###### *HG003 (NA24149), GRCh38*

ftp://ftp-trace.ncbi.nlm.nih.gov/giab/ftp/release/AshkenazimTrio/HG003_NA24149_father/NISTv3.3.2/GRCh38/

###### *HG003 (NA24149), GRCh37*

ftp://ftp-trace.ncbi.nlm.nih.gov/giab/ftp/release/AshkenazimTrio/HG003_NA24149_father/NISTv3.3.2/GRCh37/

###### *HG004 (NA24143), GRCh38*

ftp://ftp-trace.ncbi.nlm.nih.gov/giab/ftp/release/AshkenazimTrio/HG004_NA24143_mother/NISTv3.3.2/GRCh38/

###### *HG004 (NA24143), GRCh37*

ftp://ftp-trace.ncbi.nlm.nih.gov/giab/ftp/release/AshkenazimTrio/HG004_NA24143_mother/NISTv3.3.2/GRCh37/

###### *HG005 (NA24631), GRCh38*

ftp://ftp-trace.ncbi.nlm.nih.gov/giab/ftp/release/ChineseTrio/HG005_NA24631_son/NISTv3.3.2/GRCh38/

###### *HG005 (NA24631), GRCh37*

ftp://ftp-trace.ncbi.nlm.nih.gov/giab/ftp/release/ChineseTrio/HG005_NA24631_son/NISTv3.3.2/GRCh37/

#### Illumina Data

##### *HG001 (NA12878), GRCh38*

ftp://ftp-trace.ncbi.nlm.nih.gov/giab/ftp/data/NA12878/NIST_NA12878_HG001_HiSeq_300x/NHGRI_Illumina300X_novoalign_bams/HG001.GRCh38_full_plus_hs38d1_analysis_set_minus_alts.300x.bam

*# Further down-sampled to 50x using command* 'samtools view -s 0.2' *(Li, et al., 2009).*

##### *HG001 (NA12878), GRCh37*

ftp://ftp-trace.ncbi.nlm.nih.gov/giab/ftp/data/NA12878/NIST_NA12878_HG001_HiSeq_300x/NHGRI_Illumina300X_novoalign_bams/HG001.hs37d5.300x.bam

*# Further down-sampled to 50x using command* 'samtools view -s 0.2' *(Li, et al., 2009).*

##### *HG002 (NA24385), GRCh38*

ftp://ftp-trace.ncbi.nlm.nih.gov/giab/ftp/data/AshkenazimTrio/HG002_NA24385_son/NIST_HiSeq_HG002_Homogeneity-10953946/NHGRI_Illumina300X_AJtrio_novoalign_bams/HG002.GRCh38.60x.1.bam

##### *HG002 (NA24385), GRCh37*

ftp://ftp-trace.ncbi.nlm.nih.gov/giab/ftp/data/AshkenazimTrio/HG002_NA24385_son/NIST_HiSeq_HG002_Homogeneity-10953946/NHGRI_Illumina300X_AJtrio_novoalign_bams/HG002.hs37d5.60x.1.bam

##### *HG003 (NA24149), GRCh38*

ftp://ftp-trace.ncbi.nlm.nih.gov/giab/ftp/data/AshkenazimTrio/HG003_NA24149_father/NIST_HiSeq_HG003_Homogeneity-12389378/NHGRI_Illumina300X_AJtrio_novoalign_bams/HG003.GRCh38.60x.1.bam

##### *HG003 (NA24149), GRCh37*

ftp://ftp-trace.ncbi.nlm.nih.gov/giab/ftp/data/AshkenazimTrio/HG003_NA24149_father/NIST_HiSeq_HG003_Homogeneity-12389378/NHGRI_Illumina300X_AJtrio_novoalign_bams/HG003.hs37d5.60x.1.bam

##### *HG004 (NA24143), GRCh38*

ftp://ftp-trace.ncbi.nlm.nih.gov/giab/ftp/data/AshkenazimTrio/HG004_NA24143_mother/NIST_HiSeq_HG004_Homogeneity-14572558/NHGRI_Illumina300X_AJtrio_novoalign_bams/HG004.GRCh38.60x.1.bam

##### *HG004 (NA24143), GRCh37*

ftp://ftp-trace.ncbi.nlm.nih.gov/giab/ftp/data/AshkenazimTrio/HG004_NA24143_mother/NIST_HiSeq_HG004_Homogeneity-14572558/NHGRI_Illumina300X_AJtrio_novoalign_bams/HG004.hs37d5.60x.1.bam

##### *HG005 (NA24631), GRCh38*

ftp://ftp-trace.ncbi.nlm.nih.gov/giab/ftp/data/ChineseTrio/HG005_NA24631_son/HG005_NA24631_son_HiS eq_300x/NHGRI_Illumina300X_Chinesetrio_novoalign_bams/HG005.GRCh38_full_plus_hs38d1_analysis_set_minus_alts.300x.bam

*# Further down-sampled to 50x using command* ‘samtools view -s 0.2’ *(Li, et al., 2009).*

##### *HG005 (NA24631), GRCh37*

ftp://ftp-trace.ncbi.nlm.nih.gov/giab/ftp/data/ChineseTrio/HG005_NA24631_son/HG005_NA24631_son_HiSeq_300x/NHGRI_Illumina300X_Chinesetrio_novoalign_bams/HG005.hs37d5.300x.bam

*# Further down-sampled to 50x using command* ‘samtools view -s 0.2’ *(Li, et al., 2009).*

#### The reference genome

##### *GRCh38*

ftp://ftp.ncbi.nlm.nih.gov/genomes/all/GCA_000001405.15_GRCh38/seqs_for_alignment_pipelines.ucsc_ids/GCA_000001405.15_GRCh38_no_alt_plus_hs38d1_analysis_set.fna.gz

##### *GRCh37*

ftp://ftp.1000genomes.ebi.ac.uk/vol1/ftp/technical/reference/phase2_reference_assembly_sequence/hs37d5.fa.gz

#### Commands

##### *Preprocess the baseline vcf to exclude long Indels and multi-allele variants*

##### *Pre-requisitions*

### Install rtg-tools-3.7.1 *(Cleary, et al., 2014)*.
### Install vcflib (https://github.com/vcflib/vcflib).
### The baseline.vcf.gz and high-confidence-region.bed files provided by GIAB.

###### Retain only the highly confident variants in the baseline

```
rtg-tools-3.7.1/rtg vcffilter --include-bed=high-confidence-region.bed -i baseline.vcf.gz -o
baseline.inbed.vcf.gz
```

###### Remove insertions and deletions >4bp from the baseline

```
gzip -dc baseline.inbed.vcf.gz | perl -ane 'if(/^#/){print}else{if(length($F[3])==1 &&
length($F[4])==1){print}elsif(length($F[3])==1 && length($F[4])<=5){print}elsif(length($F[3])<=5 &&
length($F[4])==1){print}}' | bgziptabix baseline.inbed.withOutSV.vcf.gz
```

###### Remove multi-allele variants

```
gzip -dc baseline.inbed.withOutSV.vcf.gz | perl -ne 'if(/^#/){print}else{$a=(split)[-
1];$a=~/^(\S+?):/;if($1 eq "1/1" || $1 eq "0/1" ||  $1 eq "1/0" || $1 eq "1|1" || $1 eq "0|1" ||  $1
eq "1|0"){print}}' | bgziptabix baseline.inbed.withOutSV.noMulti.vcf.gz
```

###### Normalize the genotype in GT tag

```
pigz -dc baseline.inbed.withOutSV.noMulti.vcf.gz | perl -ane 'if(/^#/){print}else{@a=split ":",$F[-
1];$a[0]=~s/\./0/;$a[0]=~s/\|/\//;@b=split
"/",$a[0];if($b[0]>$b[1]){$a[0]="$b[1]/$b[0]"}else{$a[0]="$b[0]/$b[1]"};$F[-1]=join ":",@a; print join
"\t", @F;print "\n";}' | bgziptabix baseline.inbed.withOutSV.noMulti.normalizeGT.vcf.gz
```

#### Merge multiple samples and train model

Please refer to the demo script “dataPrepScripts/CombineMultipleGenomesForTraining.sh” in https://github.com/aquaskyline/Clairvoyante for an illustration on how to combine the training samples from different genomes and train a model for Skyhawk. Please use the “baseline.inbed.withOutSV.noMulti.normalizeGT.vcf.gz” baseline VCF file generated by the methods aforementioned for the GetTruth.py script in Clairvoyante’s model training procedures.

#### Run GATK HaplotypeCaller

```
java -Djava.io.tmpdir=/dev/shm -Xmx50G -jar GenomeAnalysisTK-3.7.jar -T HaplotypeCaller -I alignment.rg.bam -o calls.vcf.gz -R genome.fa -nct 28
# Genome.fa is either the GRCh38 or GRCh37. alignment.rg.bam is the bam files downloaded from the aforementioned links, and post-processed with `samtools addreplacerg`.
# GATK HaplotypeCaller version 3.7 was used.
```

#### Run GATK UnifiedGenotyper

```
java -Djava.io.tmpdir=/dev/shm -Xmx50G -jar GenomeAnalysisTK-3.7.jar -T UnifiedGenotyper -I alignment.rg.bam -o calls.vcf.gz -R genome.fa -nct 28
# Genome.fa is either the GRCh38 or GRCh37. alignment.rg.bam is the bam files downloaded from the aforementioned links, and post-processed with `samtools addreplacerg`.
# GATK UnifiedGenotyper version 3.7 was used.
# GATK UnifiedGenotyper by default detects only SNP variants.
```

#### Run 16GT

```
16gt/bam2snapshot -i genome.fa.index -b alignment.bam -o output/prefix
16gt/snapshotSnpcaller -i genome.fa.index -o output/prefix
perl 16gt/txt2vcf.pl output/prefix.txt sampleName genome.fa > tmp.vcf
perl 16gt/filterVCF.pl tmp.vcf dbSNP.v138.vcf.gz | bgziptabix calls.vcf.gz
# genome.fa.index was generated according to the guide in https://github.com/aquaskyline/16GT
# 16GT commit f4dddad from https://github.com/aquaskyline/16GT was used.
```

#### Identify the false positive calls

```
rtg-tools-3.7.1/rtg vcfeval -t genome.sdf -e high-confidence-region.bed -b baseline.inbed.withOutSV.noMulti.normalizeGT.vcf.gz -c calls.vcf.gz -o benchmarkingResultsFolder
# We used RTGTools for benchmarking the variant calls and extracting the false positive variants (Cleary, et al., 2014). The genome.sdf was created using the command ` rtg-tools-3.7.1/rtg format` on the genome.fa file. The high-confidence-region.bed is a part of the truth variant datasets from GIAB.
# False positive variants are in the fp.vcf.gz file in folder ‘benchmarkingResultsFolder’.
```

#### Left Align the Variants

```
gzip -dc fp.vcf.gz | vcfleftalign -r genome.fa | bgziptabix fp.leftalign.vcf.gz
```

#### Running Skyhawk

```
python validateVar.py --chkpnt_fn trainedModels/fullv3-illumina-novoalign-hg001+hg002+hg003+hg004+hg005-hg38/learningRate1e-3.epoch100.learningRate1e-4.epoch200 --ref_fn genome.fa --bam_fn alignments.bam --vcf_fn fp.leftalign.vcf.gz --thread 1 --val_fn validationResults.output
```

#### *Annotate VCF with Skyhawk Results*

```
python annotateVCF.py --vcf_fn fp.leftalign.vcf.gz --skyhawk_fn validationResults.output
--annovcf_fn validationResults.vcf
```
